# PlasmoTron: an open-source platform for automated culture of malaria parasites

**DOI:** 10.1101/241596

**Authors:** Theo Sanderson, Julian C. Rayner

## Abstract

We have created a system which allows an inexpensive opensource liquid-handling robot to automate most aspects of bloodstage malaria parasite culture. Parasites are cultured in multiwell microplates, with their details recorded in a database. Information in the database is used to generate commands for the robot to feed, monitor and passage parasite cultures. We show that the system is capable of raising cultures after transfection and then maintaining them at desired parasitaemias, facilitiating significant scale up of both routine culture and experimental genetic modification. The PlasmoTron software is available at plasmotron.org.

## Introduction

The development of a system for continuous culture of human malaria parasites *in vitro* by Trager and Jensen (1) was among the most significant methodological breakthroughs to take place in parasitology. *Plasmodium in vitro* culture is now a routine part of lab work in research labs all around the world, but it is striking how little the culture workflow in most malaria laboratories has changed since the original development of the continuous culture system. In a period of time during which the speed of computers has increased around a million-fold, the technology, and hence the limit of throughput, for routine *Plasmodium* culture is mostly unchanged. With the exception of large, short-term culture manipulations carried out for systematic drug screening (2, 3), parasites are still cultured fundamentally as they were four decades ago. This laborious process involves regular monitoring of the culture by collecting a small sample of blood and manually smearing it across a slide, which is fixed, stained and examined by microscopy. The proportion of infected cells (the parasitaemia) is visually counted, and to maintain a healthy parasitaemia the researcher then calculates the proportion of the culture which must be discarded and replaced with fresh blood in order to keep parasite growth at a sustainable level.

While this manual approach is achievable – and arguably preferable – for small numbers of cultures, it does not scale well. Beyond around twenty parallel parasite cultures, maintenance work can easily become overwhelming, and can potentially pose ergonomic hazards. Despite the existence of automated techniques for the measurement of parasitaemia (4), we are not aware of cases where this data has been used to automate the process of passaging parasite cultures. Although techniques have been developed to change media on parasites in plates in semi-automated fashion (5), this has been limited to volumes of 200 µl. In addition it has not involved a closed-loop system, in which the robot is aware of the identity of cultures on each plate, and the parasitaemia in each, and uses this data to decide how to passage parasites.

There are two major roadblocks to automation. One is the high initial investment cost for most automation platforms, with fully-fledged cell-culture automation platforms starting at $500,000. A second is that the majority of less expensive systems for general liquid-handling are designed for protocols where exactly the same steps can be performed repeatedly. These are clearly not suitable for culturing large numbers of parasite lines, where well layouts may differ widely between plates, and where differences in parasitaemia will necessitate different operations in each well of a plate. The mere concept of parasitaemia, and the need to maintain the correct ratio of two different culture components, blood and media, is unique to parasitology and so can be difficult to implement in commercial systems which were not intended for this relatively niche purpose. A potential solution is the OpenTrons system, which is considerably cheaper than any other liquid-handling system, and also very flexible, meaning that it could potentially be adapted to the unique requirements of *Plasmodium* culture.

We have used this platform to construct a system for *Plasmodium* culture which moves the responsibility for thinking about the calculations required for routine parasite maintenance from the researcher to a machine. The system is able to keep track of every aspect of large number of parasite cultures: where they are, what it expects their current parasitaemia to be, and when they were last fed. It can use this information to decide what operations need to be performed upon each culture in order to maintain a healthy population of parasites and – with the help of a researcher who supplies the plates, the media and the tips – it can perform the liquid handling functions needed to monitor, feed and passage parasites.

## Features

PlasmoTron, when used to control an OpenTrons robot ($4,000) provides the following features:

- a database containing details of all cultures, a history of all operations performed on them (feeds and passages) and their location in the rows and column of a multiwell plate.
- an automated system to feed parasites in these plates
- the ability to take an aliquot of each cuture into a measurement plate, from which parasitaemia can be read with a flow cytometer and uploaded back to the server, meaning that each culture has a record of its parasitaemia over time
- the ability to passage parasites automatically, using parasitaemia data generated from flow cytometry
- a prediction of the anticipated parasitaemia in any culture based on the most recent measurements extrapolated over time at standard growth rates

## Architecture

PlasmoTron has three key software components: a database, a web application and a robot-controller. These run on a dedicated computer and send commands to the liquid-handling robot.

### Hardware requirements

PlasmoTron is initially developed for use with an OpenTrons OT-One S Hood robot ($4,000) - equipped with a P1000 pipette and P200 8-channel pipette. These pipettes are compatible with many brands of tip, reducing consumables costs. The other requirement is a computer to run the OpenTrons robot, and host the PlasmoTron server. We used a $35 Raspberry Pi, with a touchscreen for an additional $40. The hardware set-up is shown in Fig. 1.

**Fig. 1.**
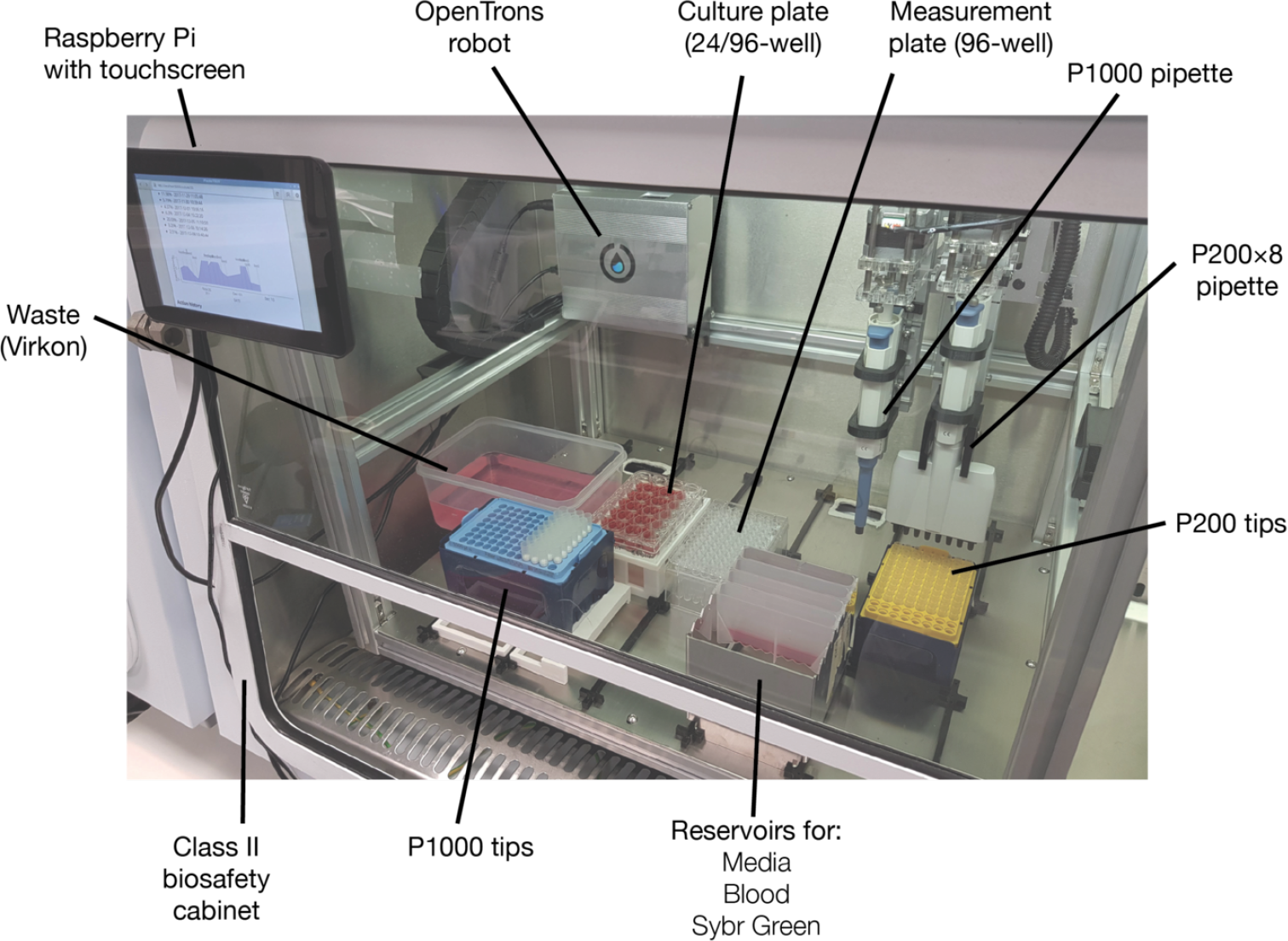
Photograph showing one hardware set up for PlasmoTron. A Raspberry Pi computer with touch screen ($75) is affixed to the outside of a biosafety cabinet. Inside the cabinet sits an OT-One Hood robot ($4,000). Parasites are cultured in 24-well or 96-well plates with 96-well plates used to collect small aliquots for parasitaemia measurement using a flow cytometer.

### Database

At the heart of the software is an SQLite3 database which includes the following tables:

- **Plates** - details of all plates, their purpose (culture or measurement) and their geometry (96-well or 24-well)
- **Cultures** - details of all cultures, their names and whether they are still under active growth
- **PlatePositions:** the positions of cultures on culture plates, and the identity of cultures on measurement plates (as well as the time when the measurement aliquot was taken)
- **Measurements:** the readings of parasitaemia from measurement plates, uploaded from a flow-cytometer. These are assigned to a certain position on a measurement plate, from which they are automatically linked to the culture from which they originated.
- **Actions:** The history of experimental actions that have been carried out on each culture (feeding, passaging, etc.)
- **CommandQueue:** raw instructions to the robot that are to be carried out or have been carried out

### Web application

The web application (Fig. 2) is a user-friendly interface to this underlying database. It permits the user to add plates to the system and populate them with cultures, and to view current parasitaemia data and upload new data. In addition it translates high-level instructions given through it’s interface (“feed plate X”, “passage plate Y to a maximum parasitaemia of 2%”) into the raw commands to the robot needed to achieve such goals, which are stored in the command queue.

**Fig. 2.**
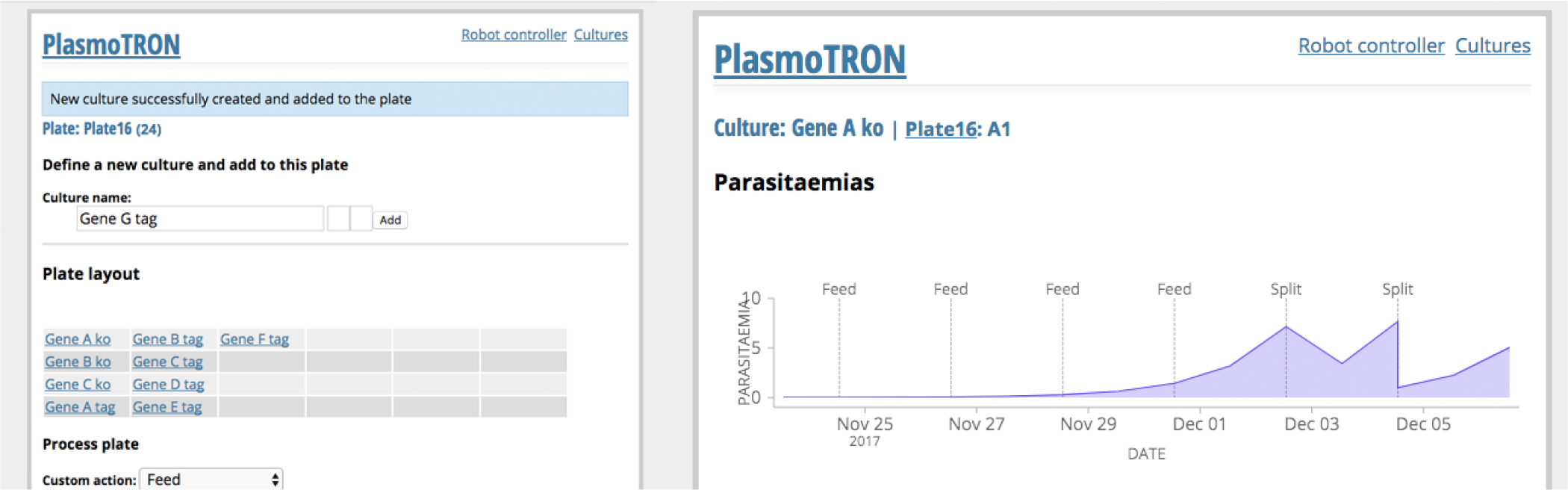
Screenshots of the PlasmoTron web application. Left: A plate is shown in the process of being populated with cultures. Right: A specific culture is shown with parasitaemia data and actions performed visualised over time.

For each culture, a graph is provided of parasitaemia over time, as well as details of every manipulation that has been performed upon it.

The application is written in Python using the Flask framework.

### Robot controller

The robot controller is a simple Python script which regularly polls the CommandQueue table in the database for instructions, and executes them by sending commands to the OpenTrons robot.

However to allow this simple system to feed back information into the database relating to the actions it has performed, some commands feature have an “onExecute” SQL statement which the robot controller runs on the database after executing them. This updates the database to reflect the changes made by the command, e.g. recording that a feed has happened, or adding a new well to a measurement plate after it is populated. In this way the database is constantly kept up to date, and if a particular operation is interrupted the system can pick up from where it left off.

The robot controller is a modular component - a different robot controller script could easily be written to control any liquid handling robot that allows a sufficient level of control.

## Methods

### Format

PlasmoTron is currently designed for culturing parasites at 1000 µl volumes in 24-well plates and 200 µl volumes in 96- well plates. In both cases, 96-well plates are used to collect aliquots for flow cytometry. However the codebase can be readily modified to almost any plate arrangement. The format of parasites within the plate can be customised arbitrarily and cultures can be archived once they are discarded. Since the database is aware of the exact makeup of the plate it is not necessary to specify for any operation how many wells need to be manipulated, or where on the plate they are located.

### Feeding

Parasites are fed by aspirating spent media from just above the layer of red blood cells settled at the bottom of each well, and subsequently replacing with fresh media. The culture is then resuspended by pipetting up and down at several different positions in the well.

### Taking an aliquot

If measurement of parasitaemia is desired then during the feeding process an aliquot of each culture is taken after fresh media has been added and the culture has been resuspended. A 20 µl sample is then taken and added to a 96-well measurement plate pre-filled with 200 µl 2× SYBR Green I per well. A record is added to the database to record the origin of each aliquot in the measurement plate. One measurement plate can be used for many culture plates, for example aliquots can be taken from four 24-well-plates and loaded onto a single 96-well measurement plate.

### Flow cytometric measurement of parasitaemia

We find that parasitaemia can be read on a flow-cytometer immediately after the measurement plate has been filled (although a short incubation might enhance sensitivity and simplify gating). We use a BD Cytoflex cytometer, but any cytometer capable of measuring parasitaemia would work well. The cytometer software is set up with a basic gating strategy which identifies single erythrocytes using FSC and SSC, and then calculates the proportion of these that are infected based on SYBR Green fluorescence, which detects DNA present in the parasite nucleus. The parasitaemia of each well in the measurement plate can then be exported directly from the CytExpert software in CSV format.

Provided the computer on which PlasmoTron is installed is networked, the web application will be accessible from anywhere on the internet (or intranet, according to network policies). The user can therefore use the browser of the flow cytometer PC to navigate to the PlasmoTron server, and upload the parasitaemia data into the record of the measurement plate within the server.

### Passaging parasites

The parasitemia data are then immediately visible on the PlasmoTron web app. Cultures can be viewed across all plates in decreasing order of parasitaemia - such a view rapidly identifies the cultures of high parasitaemia which may need to be passaged. The researcher can then load the plates containing these cultures back into PlasmoTron, specify the maximum parasitemia they want on the plate, and the system will dilute only those wells that are currently above that parasitemia, ignoring any well below the target parasitaemia. For wells above the target parasitaemia a calculated portion of culture will be discarded following resuspension. A mixture of blood and media (preprepared by the researcher) will then be added to replace the discarded culture and meet the desired parasitemia.

In the case where extreme dilutions are required for passage it is also possible to passage to a fresh plate - avoiding any issues caused by dead space.

## Results

### Applicability and reliability

We have been using PlasmoTron during its development. We have used it for routine maintenance of 48 parallel 1000 µl cultures of *P. knowlesi*, and also for bringing up *P. falciparum* transfections after electroporation. For routine maintenance, 48 cultures at 1000 µl volume could be maintained at high parasitaemia with less than an hour of operator time on Monday, Wednesday and Friday of each week. Passaging cultures to a target parasitaemia results in a range of values within a tight range of the target value (Fig. 3).

**Fig. 3.**
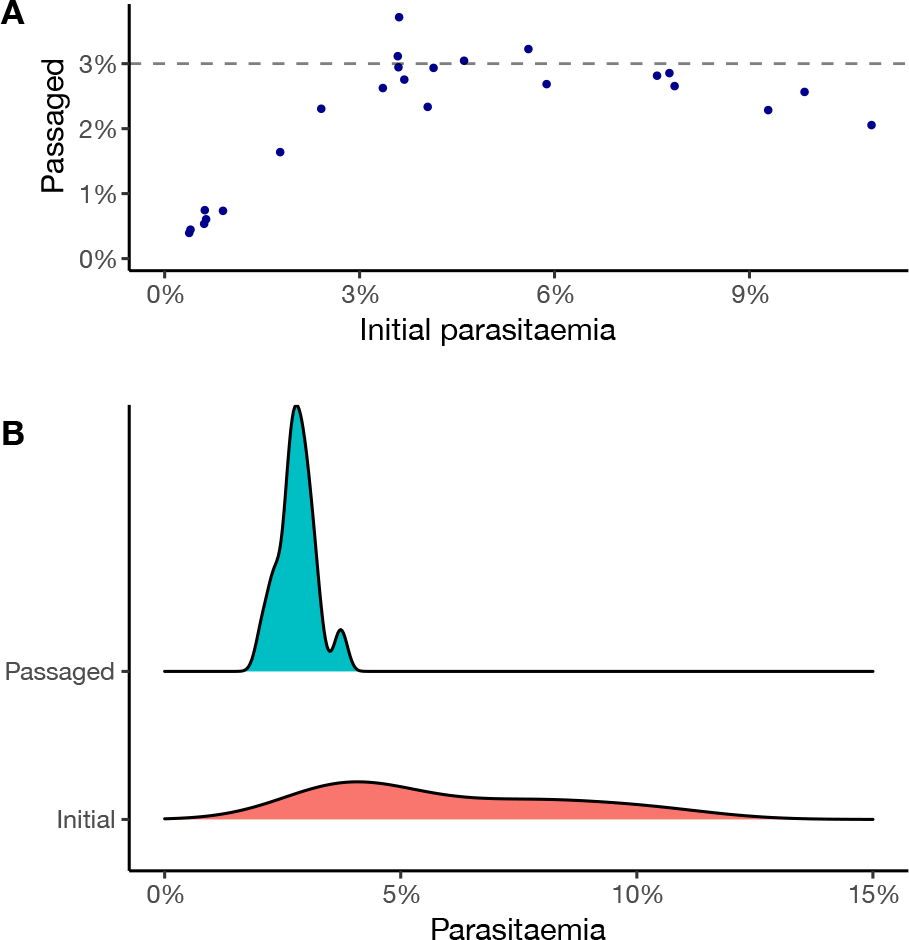
PlasmoTron accurately passages parasites to a target parasitaemia. A 24-well plate was populated with cultures of various parasitaemias from 0 to 12%. PlasmoTron was used to measure parasitaemias in all wells, and then passage the entire plate to a target parasitaemia of 3%. Final parasitaemias were again measured by flow cytometry. (A) A scatter plot of initial and final parasitaemia reveals unchanged parasitaemias below the target, whereas wells that initially had higher parasitemias had been diluted to near the target value. (B) A density plot showing only those cultures that were initially above the target parasitaemia, compared to their parasitemia after passage, demonstrates that PlasmoTron is able to reach a target parasitaemia with reasonable consistency and precision.

In the transfection experiment, parasites cultured by PlasmoTron came up at the same time as those cultured in parallel manually and there was no evidence for a reduction in the efficiency of parasite recovery caused by the use of a robotic system. No evidence of cross-contamination between wells was detected: negative wells remained negative in both manual and automated culture.

### Ease of use

Setting up the the PlasmoTron software, requires certain informatic skills (the use of the terminal for example), but once the system is installed it can be readily used by any user via the touch screen interface.

The networked nature of the robot has a number of advantages. Users can connect from any internet-enabled device, including phones and tablets, to monitor their cultures, and the robot status. From their office machines, researchers can access the robot database to view the estimated current parasitaemia of their cultures. They can also see if the robot is currently in use, to decide whether it is time to head to the laboratory. Once their plate is loaded a user can monitor the robot’s progress remotely.

## Discussion

We believe that PlasmoTron represents a significant innovation in how malaria parasites are cultured. It substantially reduces the extent to which a researcher can feel overwhelmed by their parasite cultures, and permits increases in throughput. As *Plasmodium* reverse genetic studies move to ever greater scales, such approaches will be vital.

Of course, manual culture of parasites will continue to have a place in malaria biology for the foreseeable future. A manual approach is the best way to generate the large volumes of culture needed for proteomic experiments, and some techniques such as Giemsa smears and density gradient separation are likely to be very difficult to automate. We have not yet performed any parasite synchronisation with PlasmoTron. Synchronisation is an important procedure in malaria biology, but one which often requires a centrifuge. It will be interesting in the future to attempt to expand PlasmoTron’s flow-cytometric enumeration of parasite numbers that include the distribution of different lifecycle stages, perhaps by incorporating an RNA stain (6).

Despite these limitations, for routine culture of large numbers of parallel parasite cultures, whether for high-throughput reverse genetics, selecting drug resistance, or for dilution cloning, automated approaches are likely to have great promise.

We believe that our approach reduces the cost of automating parasite culture below that of any other system. Nevertheless the approach as outlined here does require access to a flow cytometer. Lysis-based SYBR Green assays have been used to assess parasitaemia with plate-readers in similar applications and may represent a viable approach, though this requires an additional wash step (5) for robust estimation. The PlasmoTron software itself merely requires a CSV file assigning a parasitaemia to each well of the measurement plate and so should be adaptable to any high-throughput technique for assessing parasitaemia.

Automation is likely to transform the way molecular biology is carried out across diverse fields. This work provides a picture of how it may help to increase throughput for *Plasmodium* culture. The development of open-hardware approaches to integrate automated parasite incubation into a liquid-handling system is still needed, to allow a ‘set it and forget it’ approach to parasite recovery after transfection.

## Supplementary data

As part of this work we designed 3D-printed components that enable OpenTrons robots to be used in small culture hoods (Supplement 1).

## ACKNOWLEDGEMENTS

We would like to thank Mehdi Ghorbal and Rachael Coyle for acting as test-subject robot operators during system development. This preprint is formatted using a 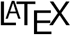 class by Ricardo Henriques.

